# Multiple-timescale dynamics, mixed mode oscillations and mixed affective states in a model of Bipolar Disorder

**DOI:** 10.1101/2022.03.22.485375

**Authors:** Efstathios Pavlidis, Fabien Campillo, Albert Goldbeter, Mathieu Desroches

## Abstract

Mixed affective states in bipolar disorder (BD) is a common psychiatric condition that occurs when symptoms of the two opposite poles coexist during an episode of mania or depression. A four-dimensional model by A. Goldbeter [27, 28] rests upon the notion that manic and depressive symptoms are produced by two competing and auto-inhibited neural networks. Some of the rich dynamics that this model can produce, include complex rhythms formed by both small-amplitude (subthreshold) and large-amplitude (suprathreshold) oscillations and could correspond to mixed bipolar states. These rhythms are commonly referred to as *mixed mode oscillations (MMOs)* and they have already been studied in many different contexts [7, 50]. In order to accurately explain these dynamics one has to apply a mathematical apparatus that makes full use of the timescale separation between variables. Here we apply the framework of multiple-timescale dynamics to the model of BD in order to understand the mathematical mechanisms underpinning the observed dynamics of changing mood. We show that the observed complex oscillations can be understood as MMOs due to a so-called *folded-node singularity*. Moreover, we explore the bifurcation structure of the system and we provide possible biological interpretations of our findings. Finally, we show the robustness of the MMOs regime to stochastic noise and we propose a minimal three-dimensional model which, with the addition of noise, exhibits similar yet purely noise-driven dynamics. The broader significance of this work is to introduce mathematical tools that could be used to analyse and potentially control future, more biologically grounded models of BD.

## 1 Introduction

### 1.1 Psychiatric Disorders from a Dynamical systems perspective

There is an ever-evolving set of paradigms taking place in Theoretical Neuroscience. Cognition, a set of computations that produces meaningful behavior, can be understood as transformations of representations within or between neural state spaces. The neural population now becomes the central implementational unit, replacing in a way the notion of neural circuits and the simple passage of messages between neurons. The so-called *state space*, that is the space of all possible states taken by the model’s variables, allows to depict the collective change of activation of the different neuronal populations. In these representation spaces, the aforementioned transformations can be described as dynamical phenomena such as equilibria (stationary states), limit cycles (robust oscillatory states), bifurcations (transitions from one type of observed state to another) [68, 4, 49]. Since psychiatric illnesses can be loosely summarised as productions of maladaptive beliefs and behavior, one could partly attribute that to disregulation of mechanisms pertaining to cognition. Furthermore, psychiatric disorders are phenomena that originate from the interplay of biophysical factors evolving on different spatial and temporal scales, while different factors can give rise to the same dynamical phenomena which, however, would result in different behavioral effects depending on the brain region where they take place [20]. All these considerations motivate us to take a new look at psychiatric illnesses through the lenses of Dynamical Systems theory, which is precisely the part of mathematical modeling that classify and analyses the possible types of states that a given time-varying system can admit for a given parameter set as well as transitions from state to state upon parameter variations. Some of the most promising approaches include the study of behavior as the outcome of a series of transient states called metastable states [40] that can be achieved through various switching mechanisms like heteroclinic channels [12] and dynamical systems with multiple timescales [38]. In the present work, we revisit a dynamical model of BD from the perspective of multiple-timescale dynamics, in order to uncover the switching mechanism that explains the metastability between mania and depression and gives rise to complex oscillations of mood. Interestingly, these complex oscillations include prolonged plateaus in the time series, which may bear relevance to mixed bipolar states or episodes (pauses) within classical bipolar oscillations from manic to depressive states and back; see Section 1.3 below for details.

### 1.2 Bipolar Disorder (BD) and models of the disease

BD is characterised by a pathological mood cyclicity between manic (extremely elevated mood) and depressive (extremely low mood) [26] episodes, interspersed with milder mood fluctuations or, in some cases, relative mood stability. Its prevalence is estimated to be within 0.3-1.5% of the total population [71]. Although small steps have been made towards the elucidation of the mechanisms of the disease, its molecular, cellular and network bases remain unknown [61, 5]. Neuronal alterations spanning all levels of description have been implicated in the pathophysiology of this complex illness [43]. Regarding the large-scale activity in BD patients’ brain, there is evidence of disturbances of the structural, functional and effective connectivity in networks responsible for affective processes, as well as those responsible for cognitive control and executive functions. These disruptions of the connectivity contribute significantly to the mood instabilities that are observed in BD [48]. In contrast, at the micro-scale level of description, the cytoarchitecture of specific brain areas bears significant changes for those patients as well. Specifically, reduction in the density of neuronal and glial cells as well as glial hypertrophy have been observed in the dorsolateral prefrontal cortex (DLPFC) of BD patients [52]. The contribution of neuronal changes, spanning different scales of the brain, to the development of BD symptoms is another factor that places Dynamical Systems theory as a key tool for the analysis of such a complex disorder.

The need to further understand the complexity of this spectrum of mental disorders resulted in various modelling attempts. In the absence of a reliable biomarker that could accurately track the evolution of this clinical entity, most of the evaluations of patients are based on their responses to questionnaires that capture in regular time periods their emotional status. In recent years, there have been a few different modeling approaches for the behavioral time series obtained from BD patients including (a) behavioral activation system models [13, 63] where altered mood stems from the dysregulation of systems governing behavioural activation or approach and (b) discrete-time random models [25]. Moreover, another category BD models is (c) biological rhythm models [14, 27, 8] in which a periodicity assumption about the mood variation of bipolar patients is implied, which could be correlated with results that show a connection between this pathological mood cyclicity and the circadian rhythms [65] or mitochondrial fluctuations [39]. Among them, the model by Goldbeter is based on the delayed interaction of two mutually inhibiting neuronal populations [27] and produces particularly interesting complex oscillatory dynamics in the BD regime. These complex oscillations, which we will show correspond to MMOs with multiple timescales, are termed “mixed state” by Goldbeter and one could also relate them to “inter-episode mood instability” as proposed by Bonsall et al. [9].

### 1.3 Mixed States in bipolar disorder and relation to MMOs in neuron models

Mixed affective states are defined as the coexistence of both manic and depressive symptoms, within a single episode of Mania or Depression. For example, a person in an episode with mixed features could be crying and having feelings of remorse while being hyperactive and experiencing racing thoughts and rapid speech. Or inversely, they could be feeling ecstatically happy and then suddenly collapse to a depressive state. In the updated version of the book *Diagnostic and Statistical Manual of Mental Disorders* [1], the term “mixed episodes” has been replaced by the specifier “mixed features” to describe low-grade symptoms of the opposite pole. The occurrence of these states is related to higher severity and worse course of illness [62], and the prevalence of these episodes is approximately 40% among bipolar patients [24, 66, 31, 44]. The pathophysiological mechanisms that lead to these mixed states include genetic susceptibility related to circadian rhythms and dopamine neurotransmission as well as disturbances in the catecholamine-acetylcholine neurotransmission balance. However, further research is necessary in order to shed light onto the precise mechanism of this condition [45]. From a dynamical systems point of view, this coexistence of manic and depressive symptoms could be expressed in the Goldbeter model through the occurrence of small-amplitude oscillations exhibited by the variables in between large-amplitude oscillations reminiscent of classical BD states.

When time series of a mathematical model are characterised by superposed rhythms consisting of both small-amplitude and large-amplitude oscillations, one typically refer to such time series as MMOs [17]. These multiple modes of oscillations are often due to the presence of multiple timescales in the underlying model, in particular in biological models. In the present work, we revisit the Goldbeter model of BD using multiple-timescale analysis since its oscillatory solutions exhibit a dynamical behaviour consistent with MMOs. Multiple-timescale analysis of mathematical models pertinent to biological application (e.g. single-neuron or neural population models) has proved very efficient in capturing complex nonlinear oscillations of MMO type; see e.g. [57, 36, 23]. However, to the best of our knowledge, this formalism has not been applied in models of psychiatric disorders, which is the main objective of the present work. It allows us to import recent developments in Mathematical and Computational Neuroscience to the young yet fast-developing field of Computational Neuropsychiatry [15, 16, 20].

From the early 1980s onward, multiple-timescale analytical and computational tools have been developed to study neural dynamics, both to analyze existing biophysical models and also to design idealized models. This classical approach of analyzing the dynamics of neuron models using so-called slow-fast theory revealed powerful enough to explain some key aspects of the *bursting* activity of neurons that was observed experimentally. In that context, when at least two fast and one slow variables were considered, the seminal work of Rinzel enabled to analyze complex bursting oscillations [55] as observed in both biophysical and idealized neuron models, for instance in the Hindmarsh-Rose model [34] or 3D versions of the FitzHugh-Nagumo [55] and of the Morris-Lecar [37] models. Namely, the shape of the full system’s limit cycle, parts of the trajectory corresponding to the slow dynamics of the system (quiescent phase of the bursting cycle), as well as parts of the trajectory corresponding to the fast dynamics of the system (active phase of the bursting cycle).

More recently, other aspects of neural electrical activity could be captured by slow-fast models with at least two slow variables, namely the possibility for spiking activity interspersed by subthreshold oscillations as observed, e.g., in some versions of the Hodgkin-Huxley model both in networks [19] and self-coupled simplified units [70]. This other type of complex oscillations correspond to MMOs and, in such context, the solution profile of the system shows a remarkable correspondence with the underlying structure of the fast subsystem and two-dimensional slow subsystem as we will show later. These complex biological rhythms modeled within the MMO framework can be compared with experiments, both at single cell level and population level; see, e.g., [7] for examples of such experimental time series. For the Bipolar Disorder model, however, experimental data are hard to be compared with the model’s output.

In the remainder of the article, we first introduce in Section 2 the Goldbeter model and analyse it using the lenses of multiple-timescale dynamics as a two slow / two fast system. Then, in Section 3, we compute and finely describe the bifurcation structure of this model with respect to two key parameters, revealing in particular a complex organisation of MMOs in parameter space along isolated curves called *isolas*. In Section 4, we consider the effect of noise on the dynamics of the model in two different ways: first, we study the robustness of MMOs to small added noise in the full 4D model; then, we derive a reduced 3D model, with only one slow variable and added noise which plays the driving role in generating noise-induced MMOs. We review our methods of analysis and computations in Section 5. Finally, in Section 6, we conclude and give a number of perspectives regarding the broader significance of the multiple-timescale modeling approach both in BD and also in other computational neuropsychiatric problems. All technical analyses of fast and slow subsystems of the model, as well as the details of noisy versions of the model, are presented in Appendix A.

## 2 The model

We consider a four-dimensional phenomenological model of bipolar disorder (BD), introduced and first analysed by Goldbeter in [27, 28]. Variables *M* and *D* correspond to the activation of neuronal populations that are responsible for manic and depressive symptoms, respectively. These variables inhibit each other. The intuition behind this inhibition comes from the two opposing poles of mood in bipolar patients; biologically, it is supported by observations of antidepressant-induced manic episodes [67]. In these cases, it seems as if the propensity to depression keeps the manic symptoms from emerging and when this propensity is decreased with antidepressants, the manic symptoms re-emerge. Of note here is that the original model by A. Goldbeter consisted only of these two variables and what is observed is the existence of two distinct stable states (bistability). The other two variables are *F*_*M*_ and *F*_*D*_ and these are intermediate factors which could be thought of as neuromodulators. One of the versions of Goldbeter’s model postulated that *F*_*M*_ and *F*_*D*_ further activate each other (cross-activation) producing, with this mechanism, bipolar oscillations. However in this work we focus on the alternative scenario, namely that the variables *F*_*M*_ and *F*_*D*_ are produced by, and also inhibit, *M* and *D*, respectively (auto-inhibition). In this way, the two intermediate factors delay the interaction between *M* and *D*. Auto-inhibition is an ubiquitous mechanism in nature, especially for neurobiological systems [18]. The model’s equations [27, 28] read

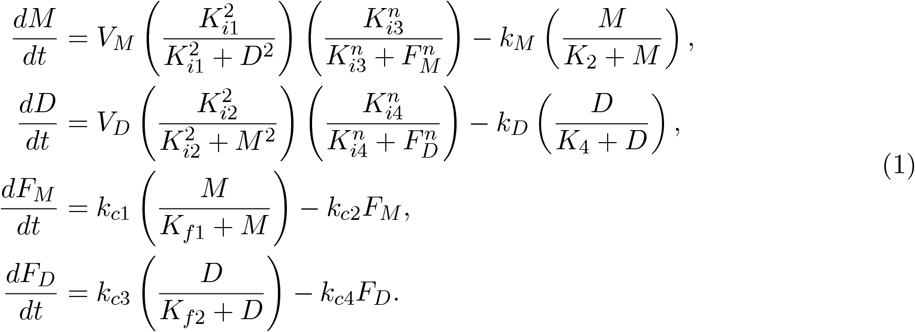

Figure 1 displays a schematic of the microcircuit underpinning this model.

**Figure 1:**
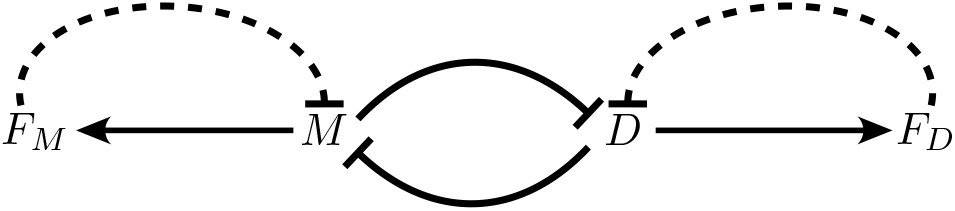
A schematic representation of the model for Bipolar Disorder that uses mutual inhibition with auto-inhibition. Adapted from [27].

### 2.1 Time series and the phase space of the model

From observing the time profile of the four variables, we obtain two important pieces of information. First, the time courses of *M* and *D* change a lot faster than *F*_*M*_ and *F*_*D*_; we reach this conclusion by observing their slopes on a sliding window; see Figure 2. Secondly, we observe that the rhythm of these variables consists an alternation between large-amplitude oscillations and small-amplitude ones. Such rhythm is often referred to as *mixed mode oscillations (MMOs)* [17]. MMOs could correspond to the mixed bipolar states that are present in about 40% of bipolar patients [44] and are characterised by the coexistence of manic and depressive symptoms during an episode of mania or depression.

**Figure 2:**
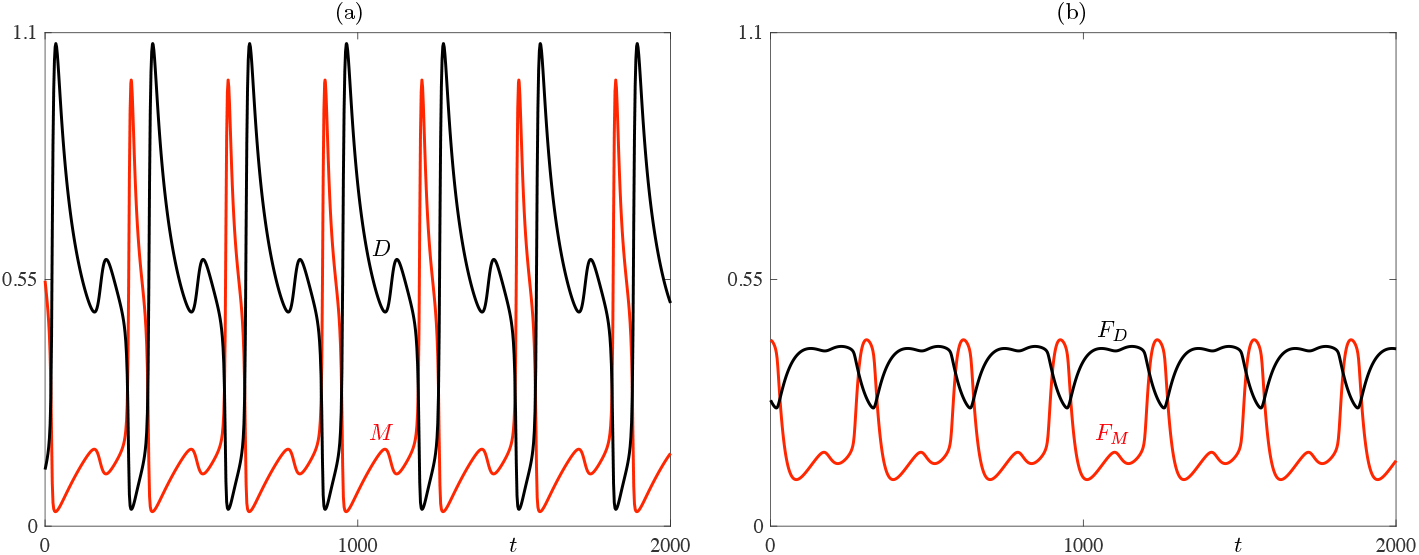
(a) Time-series of M and D displaying a MMO; (b) Time series of *F*_*M*_ and *F*_*D*_ displaying a MMO. Parameter values are as in [28, Fig.5].

Time series are key outputs of such differential equation models, and they can in some cases be compared with experimental data. However, a strong advantage of looking at a mathematical model is that we can investigate the behaviour of its solutions by plotting one variable against another one, in the so-called *phase plane*. In this context, dedicated techniques can be used in order to extract precious dynamical markers about the model. In the present case, we can see how the fast variable *M* changes with respect to the slow variables *F*_*M*_ and *F*_*D*_ along a MMO trajectory.

What we observe in Figure 3 is that there are two segments of the trajectory (in red on the figure) along which *M* changes a lot slower than the other two segments (in blue); the blue segments are “quasi-vertical” and the red ones seem to follow the lower and upper sheets of a surface represented in green which we will define below. Furthermore, we observe that there is an abrupt transition from the upper slow segment to the fast segment, while it is not the case in the transition from the lower slow segment to the fast segment of the trajectory, as there is a delay. The trajectory undergoes a turning point that corresponds to the small-amplitude oscillations mentioned above regarding the time series, which is a very important part of the MMOs. In terms of biological interpretation of this dynamical behaviour of the model, the slow segment is characterised by low manic, high depressive symptoms. In other words, one may interpret this part of the solution trajectory as a delay in the transition from depression to mania. At the core of the analysis presented in the present article is the fact that we can use classical results from multiple-timescale dynamical systems to control this delay using key parameters of the model.

**Figure 3:**
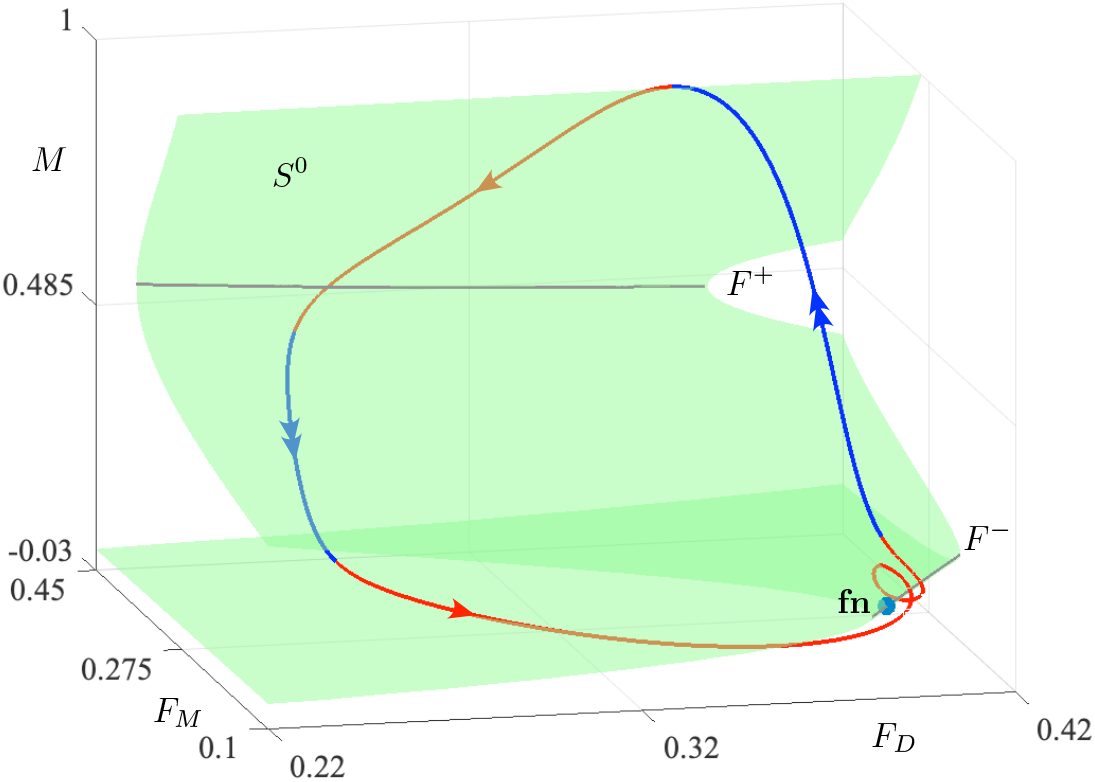
: Mixed Mode Oscillations in the Goldbeter model, as described by the system of equation in (4) (see Appendix A.1), with a mixed state close to the depressive state; parameter values correspond to Figure 5 in [28]. Also shown are the critical manifold *S*^0^, its fold curves *F* ^±^ and the folded-node singularity **fn**. Along the MMO trajectory, slow segments are highlighted in red with single arrows, and fast segments in blue with double arrows.

### 2.2 Analysing the fast dynamics

In order to exploit the different timescales present in the model and identifiable in the solutions’ time series, we first identify a small parameter responsible for the timescale separation in the equation. It turns out that, from the original parameter set considered in [27, 28], parameters 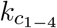 are all small and of the same order, therefore we will introduce a small parameter *ε* equal to 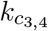, which will automatically give that 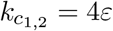 is still small.

This timescale ratio parameter *ε* plays a central role in slow-fast systems. First and foremost, it allows to identify two fast variables (*M* and *D*) and two slow variables (*F*_*M*_ and *F*_*D*_) in this BD model. When *ε* multiplies the right-hand side of the slow equations in system (4) then the system is said to be written in the *fast-time parametrization*, with the *fast time t*, in which equations (1) take the form

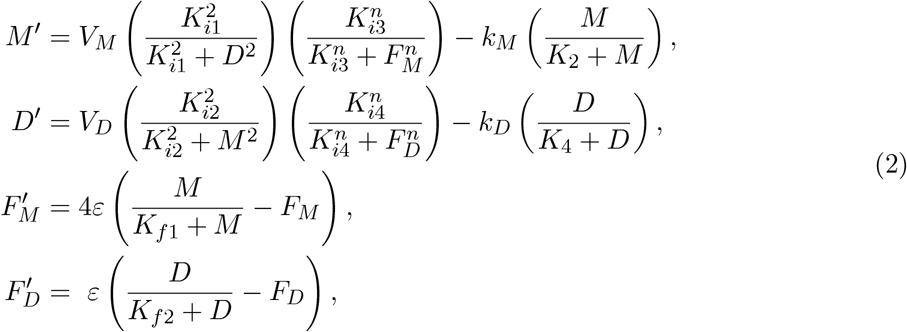

where the prime denotes the derivative with respect to the fast time *t*. If we want to explore the fast segments of the trajectory, i.e., the fast dynamics of the system, we can consider the limit of the fast-time system equations (4) when *ε* tends to 0. This is perfectly possible and legitimate in a mathematical model even though it is impossible in an experimental context. However, it still unveils important elements that one can use to decipher the complex dynamics of the full system. Essentially, in this way we freeze the dynamics of the slow variables and consider them as parameters. As a result, we get the so-called *fast subsystem* which gives an approximation of the fast dynamics of the full system.

If we solve the system of equations *M′* = 0 and *D′* = 0, we can find the set of equilibria of the fast subsystem. By plotting them in a three-dimensional phase-space projection, for instance onto (*F*_*M*_, *F*_*D*_, *M*), we obtain a surface called the *critical manifold S*^0^ of the system; this is the green surface shown in Figure 3. Superimposing the MMO trajectory of the full system onto the critical manifold, that is, onto the set of equilibria of the fast subsystem, we can see that there is a remarkable fit. This confirms that our original assumption of a two fast/two slow variable system was correct. Moreover, superimposing these solutions of the full system onto the bifurcation diagram of the fast subsystem with respect to the “frozen” slow variables proves very pertinent: the full system solution follows slowly branches of attractors of the fast subsystem and transitions between different phases appear near bifurcation points of the fast subsystem. In the present context, we have two slow variables so two main parameters in the fast subsystem; hence, important transitions along an MMO trajectory occur near lines of bifurcations of the fast subsystem, namely lines of fold bifurcation, which geometrically correspond to the two fold lines *F* ^±^ of the critical manifold; see also Appendix A.1.

Given that the system also has two slow variables, the slow singular limit at *ε* = 0 also contains valuable information, in particular about the mixed-state dynamics, as explained next. For more details on the methodology, see Appendix A.2.

### 2.3 Analysing the slow dynamics

We now turn our focus to the study of the slow segments of the trajectory, i.e., the slow dynamics of the system. For that, we can rewrite the system in *slow-time parametrisation* (7) (see Appendix A.2) by introducing the so-called *slow time τ* = *tε*. Here we use an overdot to denote differentiation with respect to *τ*:

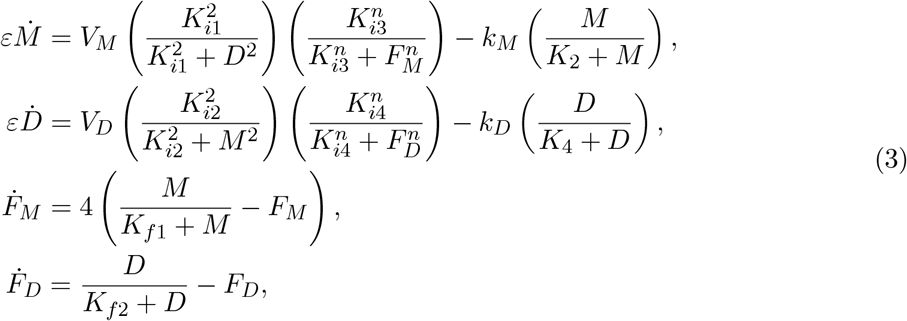

where the over dot denotes the derivative with respect to the slow time *τ*. In this case, if we take the abstract limit *ε* = 0, the resulting system is called the *slow subsystem* and it approximates the slow dynamics of the full system. It consists of two differential equations that are constrained by two algebraic equations which we can write in vector form {**f** = 0}, where **f** is a complicated nonlinear function of all the variables.

This equation {**f** = 0} is the one that we used to plot the critical manifold. In other words, the slow subsystem solution trajectories are forced to evolve on the critical manifold when *ε* = 0. Thus, the critical manifold forms both the set of equilibria of the fast subsystem, as we saw earlier, and the phase space of the slow subsystem. Because this algebraic equation {**f** = 0} has to hold true for all times *τ*, we can differentiate it with respect to time *τ*. The resulting formula is given in (9).

When the denominator vanishes and the numerator does not, the slow subsystem is not defined, and this happens along the fold curves of the critical manifold. There are, however, special points where the numerator also vanishes by having a zero of the same order as the denominator. In such cases, the slow subsystem is well defined. These points are called *folded singularities*, they correspond to turning points of the slow flow and are responsible for the aforementioned delayed transition from depression to mania. Note that in some cases, folded singularities can have an interplay with a *delayed Hopf bifurcation* and the delayed transition and accompanying small oscillations can also be related to the presence of equilibria with complex eigenvalues in the fast subsystem; see e.g. [17]. Details about the derivation and analysis of the slow flow are given in the Appendix section A.2.

By changing the value of parameter *K*_*f*1_, one can find by direct simulation another type of MMO trajectory, where the delayed transition now occurs at the transition from mania to depression; see Figure 4. Here as well, the length of the delay (i.e. the duration of the mixed state) can be controlled by adjusting the parameter *K*_*f*1_. Another folded singularity (of the same node type) is also responsible for the MMOs whose intermediate state is closer to mania (higher values of *K*_*f*1_) and located on the upper fold of the critical manifold and shown in Figure 4.

**Figure 4:**
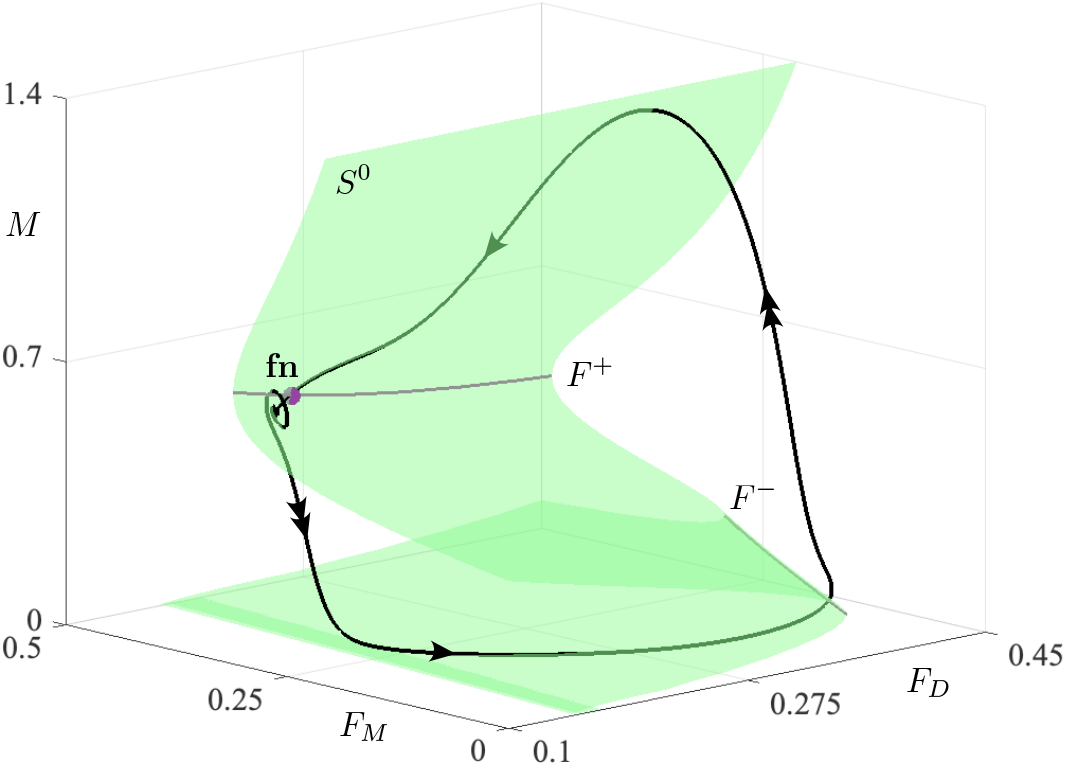
Mixed Mode Oscillations in the Goldbeter model (4) with a mixed state close to the manic state; parameter values as in Fig. 3 except for *K*_*f*1_ = 1.293. Also shown are the critical manifold *S*^0^, its fold curves *F* ^±^ and the folded-node singularity **fn**. Along the MMO trajectory, slow segments are highlighted by single arrows, and fast segments with double arrows.

The important conclusion is that the aforementioned delay in the transition from the slow to the fast segment in the MMO trajectory corresponds to a delay in the transition from depression to mania, or in the opposite transition depending on *K*_*f*1_, and such mixed bipolar states can be attributed to the existence of a folded singularity. Furthermore, by changing the values of system parameters, we can control the duration of this delay, which emphasizes the relative importance of a mixed state within a bipolar disorder episode. We therefore need to study the various types of solutions that the model admits depending on the value of a key parameter like *K*_*f*1_ and this is the purpose of the next section.

## 3 Bifurcation analysis of the system

### 3.1 Exploring the bifurcation structure with respect to *K*_*f*1_

Bifurcation analysis is a common tool in the study of non-linear dynamical systems, also in the context of neuroscientific problems (e.g. [11]), as it enables to explore how small changes in the values of the system’s parameters affect the system’s regime. Parameter *K*_*f*1_, and symmetrically parameter *K*_*f*2_, play an important role for the model. In this model, the rates of production of *F*_*M*_ and *F*_*D*_ take the form of Michaelis-Menten like functions and therefore they display saturation and reach a plateau at large values of M and D. The constants *K*_*f*1_ and *K*_*f*2_ reflect the levels of *M* or *D* achieving 50% of the maximum rates of production of *F*_*M*_ or *F*_*D*_, respectively. In other words, the rates of synthesis of *F*_*M*_ and *F*_*D*_ saturate and reach a plateau value for sufficiently low values of *K*_*f*1_ and *K*_*f*2_, while they become linear at large values of these constants. We now explore how variations of parameter *K*_*f*1_ affect the dynamical regime of the system. The bifurcation diagram of the full system with respect to *K*_*f*1_ is presented in Figure 5.

**Figure 5:**
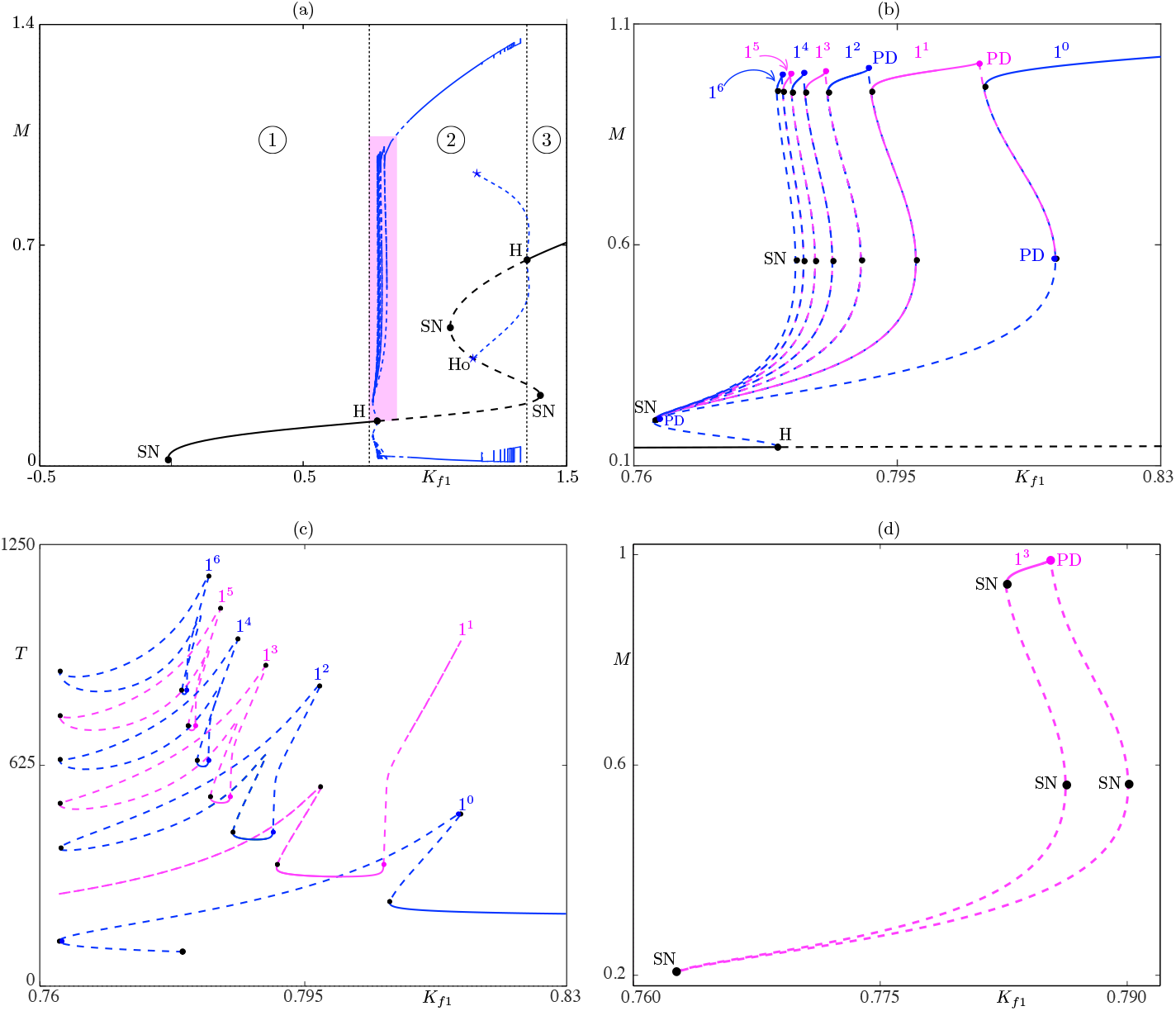
(a) Bifurcation diagram with respect to the parameter *K*_*f*1_; (b) Zoomed view of panel (a) (colored rectangle) highlighting 6 isolas MMOs corresponding to solutions with profile 1^1^ − 1^6^, respectively. The label 1^*s*^ refers to MMO with 1 large-amplitude and *s* small-amplitude oscillations per period. By extension, we call 1^0^ standard BD oscillations, with only 1 frequency so no small-amplitude oscillations. (c) Period along all computed branches of periodic solutions. (d) The isola of 1^3^ MMOs shown alone so as to highlight its geometry and the fact that it is closed in parameter space. In all panels the solid lines correspond to stable equilibria, whereas the dashed lines correspond to unstable equilibria. SN: saddle-node bifurcation, H: Hopf bifurcation, Ho: homoclinic bifurcation, PD: period-doubling bifurcation.

It appears that there are three distinct areas of interest. For low values of *K*_*f*1_, the system is in a steady state that is characterised by low levels of mania (and high levels of depression). Thus, region 1 corresponds to Depression. As we increase *K*_*f*1_ there is a Hopf Bifurcation that gives rise to a family of limit cycles. Now the system is in an oscillatory regime between manic and depressive states, hence region 2 corresponds to bipolar disorder. Lastly, if we increase further the value of *K*_*f*1_ then the system settles in a different steady state characterised by high manic (and low depressive) symptoms, which means that region 3 corresponds to mania. All 3 regions are highlighted in Figure 5 (a).

Moreover, for values of *K*_*f*1_ at the transition from region 1 to region 2, we can observe in the corresponding time-series, the kind of rhythms that were mentioned earlier, consisting of large- and small-amplitude oscillations (MMOs). These MMOs have an intermediate state closer to depression (*D* reaching higher values than *M*). The family of MMOs that is located the closest to the branch of limit cycles born at the Hopf Bifurcation consists of one large-amplitude and one small-amplitude oscillation and it is denoted as 1^1^. The second MMO branch corresponds to 1 large and 2 small oscillations (1^2^), and so on. In the parameter space, most of them are organised along closed isolated branches typically referred to as *isolas*; a few such isolas are shown in details in Figure 5 (a) and (c), and one specific isola is shown in panel (d) to showcase its closed geometry in parameter space. However, this is not the case for the first branch of MMOs (1^1^). Numerical evidence supports the fact that this branch of 1^1^ MMOs starts and ends through a period-doubling bifurcation from the main branch of limit cycles, the one that is born at the Hopf bifurcation. We provide this numerical evidence in Figure 5 (c) where we plot all MMO branches with the period as solution measure, and indeed the beginning and end of the 1^1^ branch correspond to a doubling of the period with respect to the corresponding parts of the 1^0^ branch. This is quite typical of slow-fast systems with MMOs due a folded node as studied in, e.g. [17], specifically figure 19 which was made for a prototypical model of MMO dynamics due to folded node (the so-called Koper model). Thus, what we observe in the Goldbeter model is consistent with this scenario.

MMOs with more small-amplitude oscillations exist on isolas and we show five such isolas: MMOs of type 1^2^ to MMOs of type 1^6^. Essentially, we can control the number of small oscillations by changing, for instance, the parameter *K*_*f*1_. Note that MMOs due to the presence of a folded-node singularity are very much related to *canard solutions*, known to organise, in this context, the transition between different profiles of MMOs upon parameter (e.g. *K*_*f*1_) variation [17]. By decreasing the parameter’s value we can increase the period of the MMO, hence the delay of the transition from the slow to the fast dynamics of the system and by extension the transition from the depressive to the manic state; see Figure 6 for an illustration where the number of small oscillations (hence the amount of delay to the transition to mania) progressively increases as *K*_*f*1_ is decreased in the MMO regime. For values of *K*_*f*1_ at the transition from region 2 to region 3, we can observe MMOs whose intermediate state is closer to mania. In parameter space, these families of MMOs are organised along open branches bounded by homoclinic bifurcations on both sides, which in particular indicates that the duration of the mixed state can become very large. In the limit, we observe a plateau instead of small oscillations, interspersed with abrupt spikes, or episodes, of mania.

**Figure 6:**
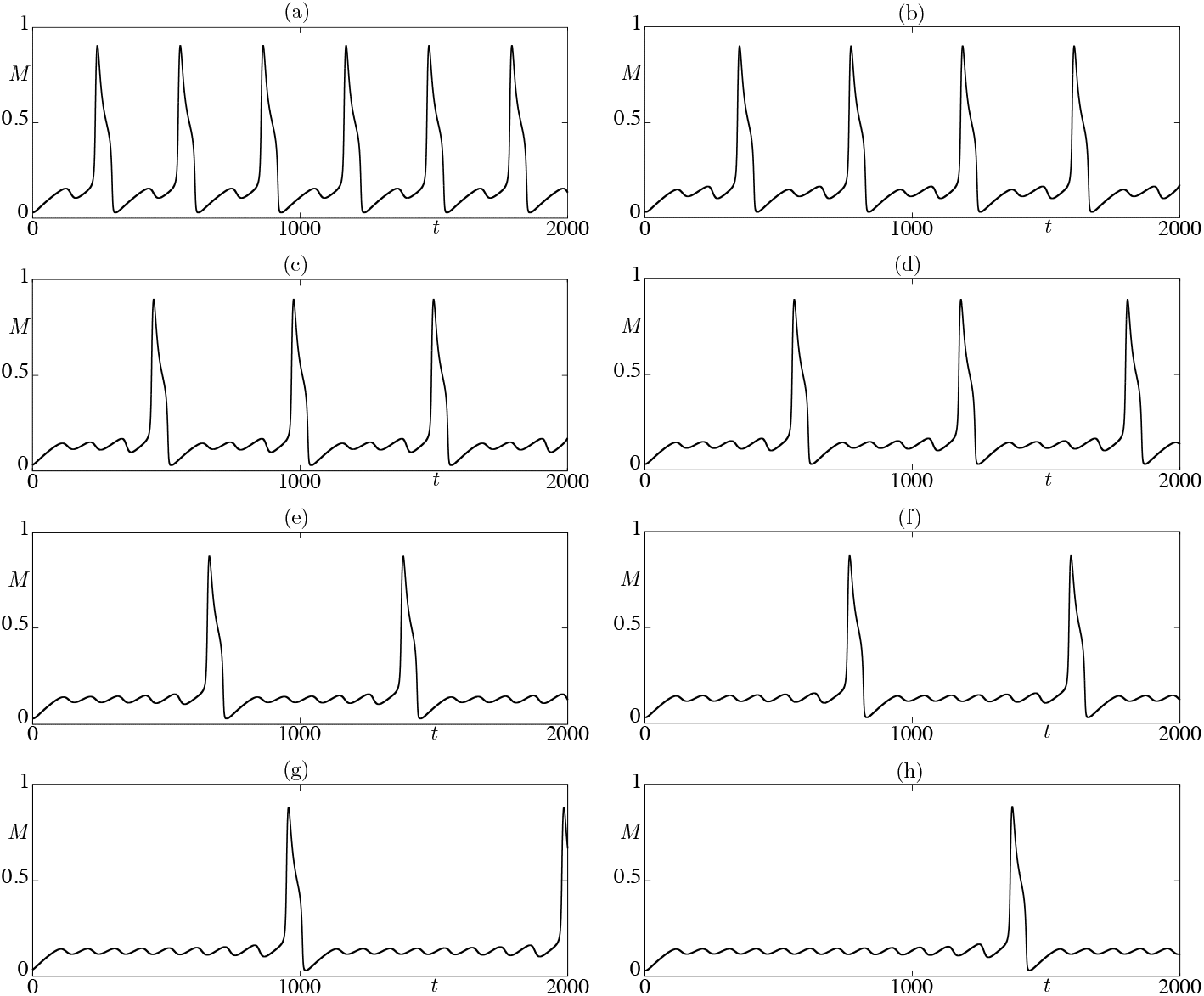
Time series for variable *M* of stable MMO solutions for various values of *K*_*f*1_, illustrating the fact that the effect of varying this parameter is to create more and more small-amplitude oscillations in between the large-amplitude oscillation displayed per period. Values of *K*_*f*1_ and MMO profiles are: (a) *K*_*f*1_ = 0.8 with a 1^1^ MMO; (b) *K*_*f*1_ = 0.79 with a 1^2^ MMO; (c) *K*_*f*1_ = 0.785 with a 1^3^ MMO; (d) *K*_*f*1_ = 0.782 with a 1^4^ MMO; (e) *K*_*f*1_ = 0.78 with a 1^5^ MMO; (f) *K*_*f*1_ = 0.779 with a 1^6^ MMO; (g) *K*_*f*1_ = 0.778 with a 1^8^ MMO; (h) *K*_*f*1_ = 0.777 with a 1^12^ MMO.

It is possible that the change of other parameters has a similar effect too. An interesting connection can be made with the bifurcation diagram of the DRS (an auxiliary system that was used for the analysis of the slow dynamics of the system; see Appendix A.2 for details) with respect to the parameter *K*_*f*1_, as illustrated in Figure 7. We observe that there are two transcritical bifurcations (denoted by T), which correspond to the birth and death of the MMO regime in the full system and the values of the parameter are similar to the ones that correspond to the MMO regimes that were mentioned before. Finally, claims of chaotic patterns of mood variation in bipolar disorder have been the focus of scientific efforts since the 1990s. Gottschalk et al. [32] used time series analysis to show that self-reported mood in patients with bipolar disorder can be characterised as a process evolving in a low-dimensional chaotic regime. More recent studies show that these patterns could be reinforced by stronger interactions between negative affective states [72]. In accordance with these observations, the present BD model can exhibit a highly irregular non-periodic behavior of chaotic oscillations which is very sensitive to even small perturbations of the initial conditions of the system and thus very hard to predict. These dynamics are common for systems that produce MMOs as shown in multiple studies [3, 41]. For a narrow range of values of parameter *K*_*f*1_, the system undergoes successive transitions to limit cycles with a period that doubles each time, known as *period-doubling cascade*. This is one of the known mechanisms of creation of deterministic chaos and it often appears in the MMO regime of multiple-timescale dynamical systems. The integration of the system for a long simulation time can give rise to a solution trajectory that is consistent with the existence of an underlying chaotic attractor because we can observe a significant (deterministic) variation of the trajectory which is, however, confined in a certain “volume” of the state space; see Figure 8.

**Figure 7:**
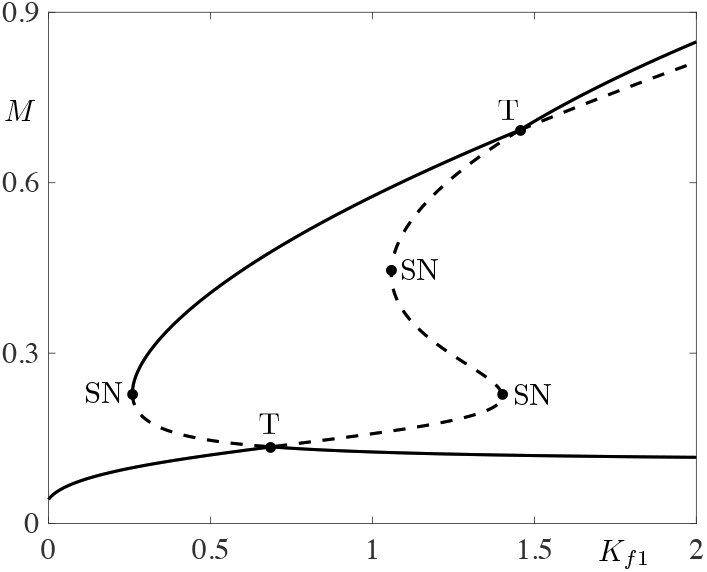
Bifurcation diagram of the Desingularized Reduced System (DRS) with respect to parameter *K*_*f*1_. The structure of this diagram reveals the presence of two types of equilibria of the DRS, namely folded singularities (which are not equilibria of the true slow subsystem) and true singularities (which are also equilibria of the slow subsystem). Both types of singularities meet at transcritical bifurcation points *T*_*i*_. In the present context, such bifurcations correspond to the event where a folded node loses stability to become a folded saddle and a (true) saddle becomes a (true) node. Hence, this event marks the boundaries of the MMO regimes.

**Figure 8:**
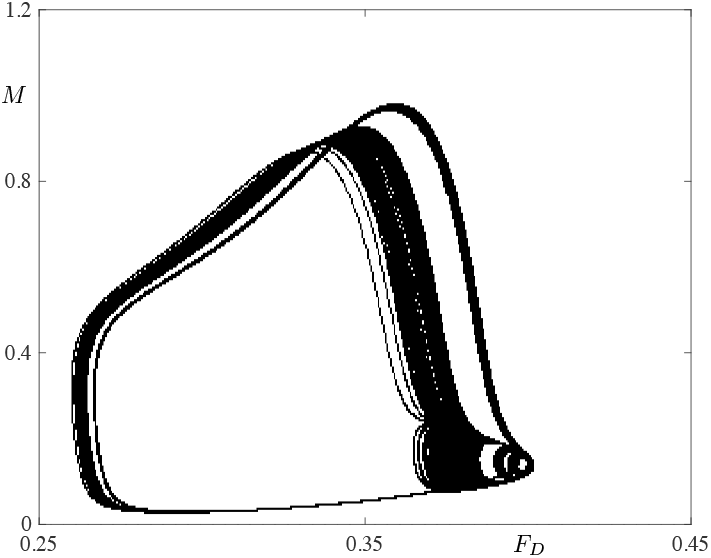
Chaotic attractor in system (4) obtained by direct simulation for *K*_*f*1_ = 0.78065832.

### 3.2 Two-parameter bifurcation analysis with respect to *K*_*f*1_ and *V*_*M*_

Another important parameter that plays a crucial role in the model of bipolar disorder is the ratio *θ* of the parameters *V*_*D*_ and 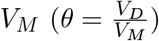. Parameters *V*_*D*_ and *V*_*M*_ dictate the maximum rate of increase of the propensity to depression and mania respectively, which are correlated with the activation of the two opposing neuronal populations D and M. An original formulation of the model which produced bistability between mania and depression was based on the increase of *θ* so that the transition from mania to depression can be achieved and vice-versa. The oscillatory behavior of the model was created when this ratio *θ* was coupled with the two main variables M and D through the equations of the intermediate factors *F*_*M*_ and *F*_*D*_ [28, 27]. Thus, switching between states in this model is strongly related to the *θ* ratio. In a way *θ* expresses the end result of various proposed triggers of the mood swings in Bipolar Disorder. These triggers can be pharmacological factors, as in the case of anti-depressant treatment-induced switching to a hypomanic or manic state, or environmental factors such as a disturbed sleep pattern, *social rhythm disruption (SRD) (social zeitgeber)* [33] due to adverse life events, or the changing of seasons [73]. Alternatively, the continuous accumulation of a multitude of risk factors, as those mentioned above, could push the person to reach a rather sudden shift (catastrophe or tipping point) from one mental state or mood to another [47, 58]. By definition of *θ*, changing *θ* for a given value of *V*_*D*_ is similar to changing the parameter *V*_*M*_. We therefore investigated numerically the change of the dynamical regime of the system with regard to the values of both *V*_*M*_ and *K*_*f*1_. The result is shown in Figure 9. The black line depicts a family of Hopf Bifurcations (HB). The red line depicts a family of saddle-node bifurcations (SN). For descending values of *V*_*M*_ we observe that the two branches of the family of SNs meet at a cusp (*V*_*M*_ ≈ 0.9) after which there are no more saddle nodes. If we want to visualise better this two-parameter bifurcation diagram, we can imagine that for *V*_*M*_ = 1 a snapshot of the bifurcations of the system would be the same as the bifurcation diagram for *K*_*f*1_ in Figure 5, with two saddle-nodes and two HBs. The important information that this bifurcation diagram conveys is the range of values of *K*_*f*1_ and *V*_*M*_ for which the system is in an oscillatory regime (bipolar disorder). The oscillations are attributed to limit cycles that are created by the HBs. And these values lie within the black curve of the HBs. The other important information is that for values of *V*_*M*_ below the codimension-two cusp point, we do not have the possibility of the two distinct states Depression and Mania that we saw before (region 1 and region 3 in Figure 5), because in the parameter space these states appear as branches of stable equilibria after the SN bifurcations only for values of *V*_*M*_ higher than the value at the cusp.

**Figure 9:**
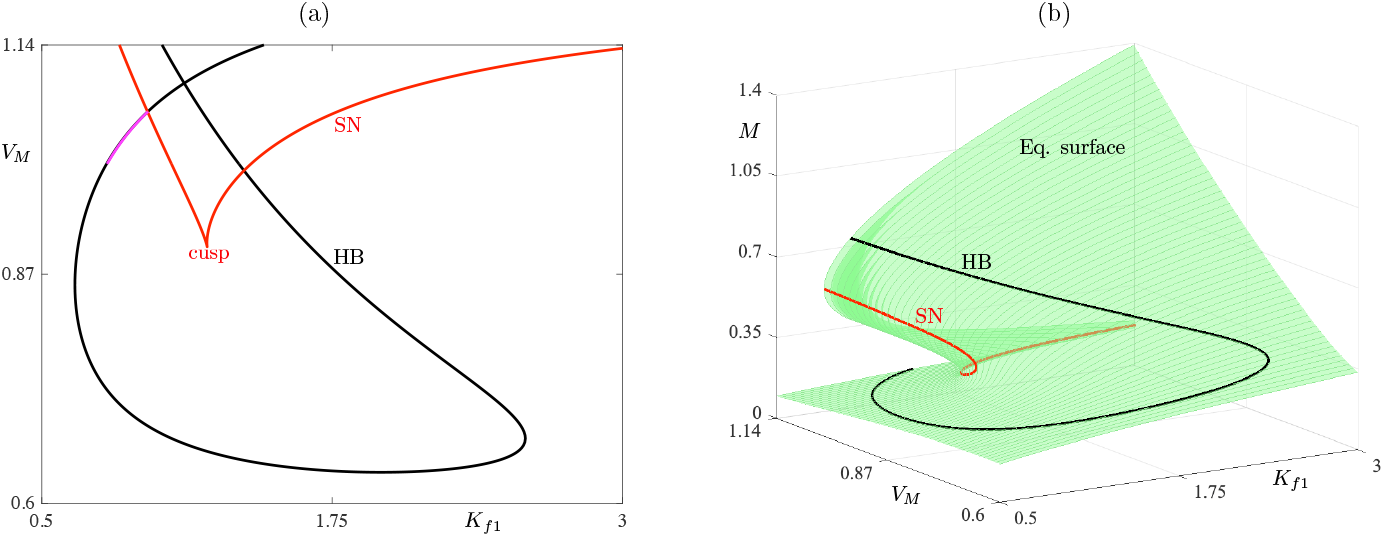
(a) Two-parameter bifurcation diagram with respect to *V*_*M*_ and *K*_*f*1_. The black line corresponds to a family of Hopf Bifurcations (HB). The purple segment of the black line corresponds to the “isolas” that were mentioned earlier. The red line corresponds to a family of saddle-node bifurcations (SN). (b) The same diagram plotted in the 3D space (*K*_*f*1_, *V*_*M*_, *M*) space and superimposed on the equilibrium manifold of the system, that is, the zero set of the right-hand side of system (1).

## 4 Robustness of deterministic MMOs to noise and noise-induced MMOs

The existence of stochastic noise is ubiquitous in biological systems and thus an important factor contributing to the emergence of psychiatric disorders [56, 51]. Hence, taking into account the effects of noise in this multi-time scale model of bipolar disorder could contribute to a more accurate explanation of real data of bipolar patients’ mood changes that are susceptible to random fluctuations. Firstly, we investigate the effect of noise on this system of ODEs. If we add a noise term (a standard Wiener process multiplied by a small parameter of the order *ε*) we observe from the time-series that the system, in general, retains the structure of MMOs. We indeed show in Figure 10 that MMOs are robust to small-amplitude noise added to the fast equations; see Appendix A.3. However, part of the small amplitude oscillations becomes less distinguishable from the random fluctuations of the values of the fast variables. Of course, for large values of the noise term this structure is lost. The relation between the noise level relative to the timescale ratio parameter plays a significant role as shown in previous works [6, 60]; see also [64] for another study investigating the effect of noise on complex oscillations with multiple timescales. A similar exploration of the effects of noise on the model’s oscillatory regime has been made by A. Goldbeter in [28].

**Figure 10:**
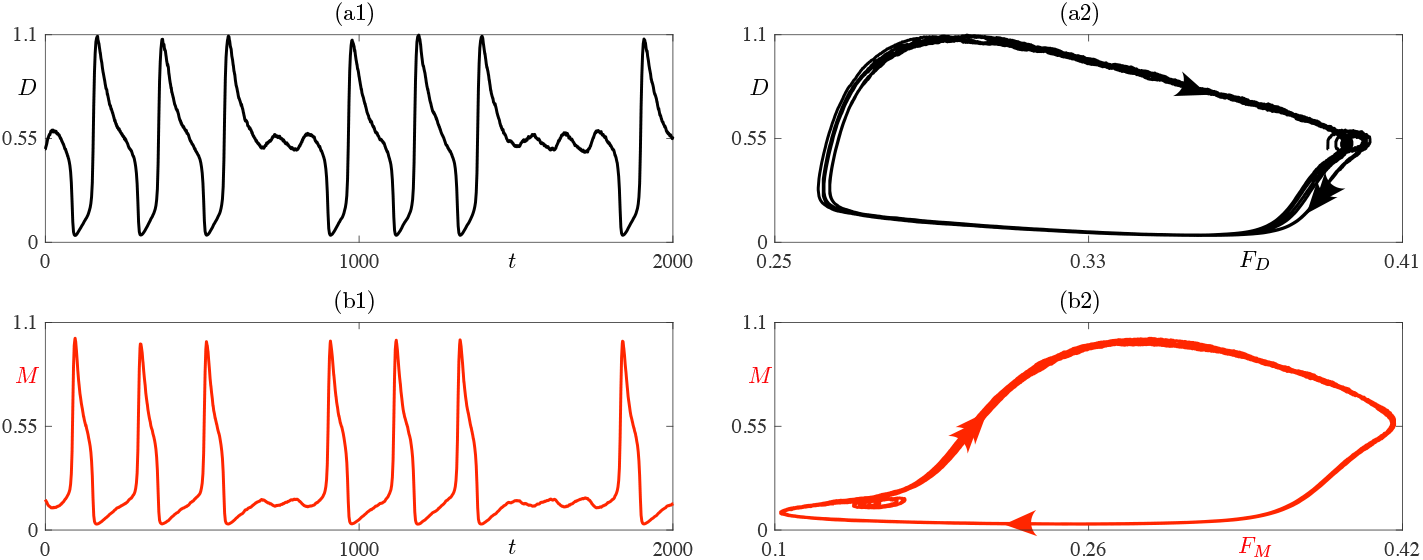
Simulation of system (4) with added Gaussian noise in the fast equations, that is, equations (13). Panels (a1)-(b1) show the time series of variables *D* and *M*, respectively. Panels (a2)-(b2) display the phase-plane projections onto the (*F*_*D*_, *D*) and onto the (*F*_*M*_, *M*) planes, respectively.

Furthermore, it is established that the MMOs can be created with various mechanisms by structurally different systems (noise induced, delay bifurcations, etc) and there is previous work that proposes ways to identify this mechanism [10]. One of the ways for a system to produce MMOs is with the addition of stochastic noise. Although in a deterministic setting the minimal requirement for MMOs is the existence of one fast and two slow variables, we can create MMO dynamics through a different approach that needs only one slow variable. To this end, we removed the slow variable *F*_*D*_ (now considered as a fixed parameter) and introduced a noise term *B* (standard Wiener process) in the remaining slow variable *F*_*M*_. This produces MMOs that are purely noise-induced, as seen from the time-series and the trajectories in the phase plane in Figure 11. Essentially the slow perturbations that lead to this peculiar periodic solution that we described as MMOs are driven by the stochastic noise instead of the interplay of the two slow variables. The important benefit of this attempt is twofold. In computational modeling we attempt in general to provide the minimal model that serves our purposes, (in our case reproduce the behavioural time-series, obtained by bipolar patients) driven by the principle of parsimony i.e simpler models and theories are preferable because they tend to be more easily falsifiable. Adding to that, mood is a phenomenon affected by the stochasticity of biological processes as well as by random life events. This attribute of mood can be captured by the model with the addition of the noise term. In other words there is a clear interpretation of the stochastic drive of the periodic solutions, namely the induction of the mixed bipolar states by random environmental factors or events. For the formulation of the respective systems of stochastic differential equations, see Appendix A.4.

**Figure 11:**
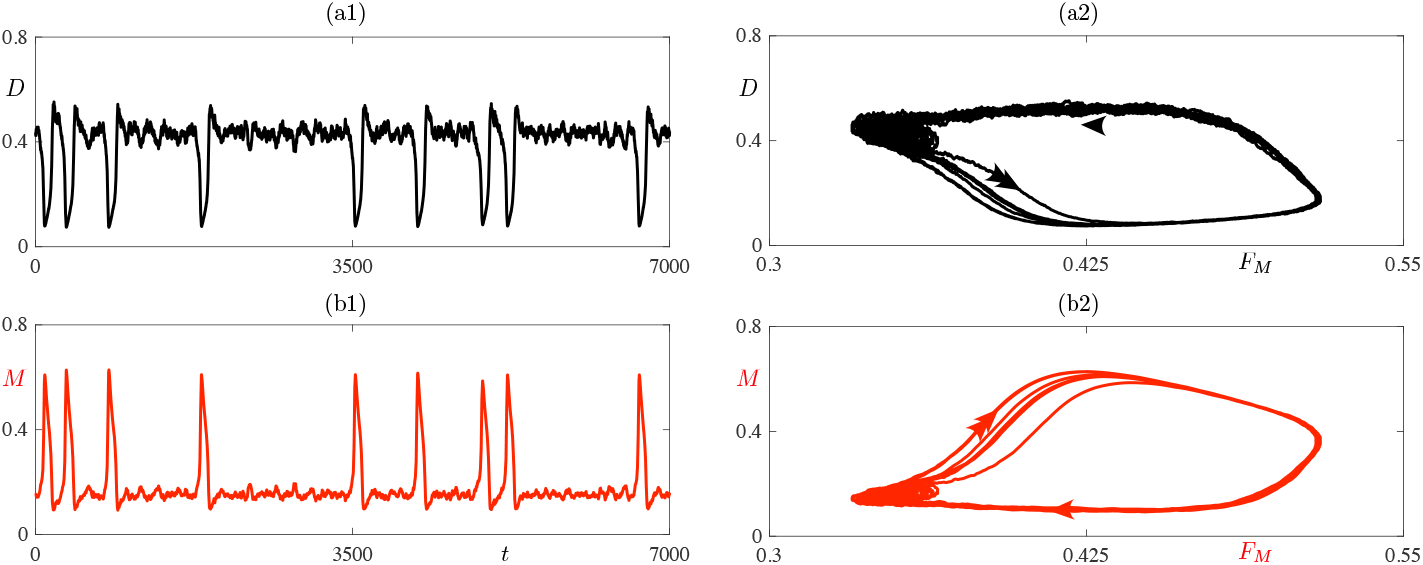
Noise-induced MMOs in a 3D reduced version of the Goldbeter model with added Gaussian noise in the slow variable; see equations (14). Panels (a1)-(b1) show the time series of variables *D* and *M*, respectively. Panels (a2)-(b2) display the phase-plane projections onto the (*F*_*D*_, *D*) and onto the (*F*_*M*_, *M*) planes, respectively.

## 5 Methods

The main methods that we have employed in order to revisit the BD model by Goldbeter are both of theoretical (slow-fast analysis, bifurcation theory) and computational (numerical solutions of ODEs and numerical continuation) nature. For the simulations of the trajectories of the system on the phase planes we used the XPPAUT software and the numerical scheme that we used was Runge-Kutta order 4. For the construction of the bifurcation diagrams, we used the numerical continuation algorithm of the XPPAUT software [22]. Numerical continuation is a standard method of computing families of attractors (e.g. stable equilibria, stable limit cycles) as well as repellor (stationary or periodic) and bifurcation points marking changes of stability along such families. On top of these, other functions are implemented which can detect bifurcation points of nonlinear systems i.e. changes of the dynamics of a dynamical system within the neighborhood of a specific parameter value. For the 3D plots in the phase-space with the critical manifold and one trajectory, we used the matlab environment. The parameter values used for the Bipolar Disorder Model were given in [27, 28]: *V*_*M*_ = 1, *θ* = 1.2, *V*_*D*_ = *θ V*_*M*_, *n* = 1, *K*_2_ = *K*_4_ = 0.5, *K*_*i*1_ = 0.33, *K*_*i*2_ = 0.35, *K*_*i*3_ = 0.6, *K*_*i*4_ = 0.4, *k*_*M*_ = *k*_*D*_ = 1, *k*_*c*1_ = *k*_*c*2_ = 0.04, *k*_*c*3_ = *k*_*c*4_ = 0.01, *K*_*f*1_ = *K*_*f*2_ = 0.8. Initial conditions: *M* = 0.161, *D* = 0.495, *F*_*M*_ = 0.165, *F*_*D*_ = 0.391. All the scripts are available upon request and will be uploaded onto ModelDB ^1^ upon publication of the present work.

## 6 Summary and Discussion

The slow-fast theory for MMO due to a folded node is rather recent (it was developed from the 1990s onward) and it provides a framework to study certain types of complex oscillations with both small-amplitude and large-amplitude components. Such complex oscillatory patterns are ubiquitous in biological signals and multiple-timescale MMO theory has proven very useful in many neural systems, both at single-cell and population level. Slow-fast theory is especially relevant to Bipolar Disorder. There is an accumulation of evidence related to the topological features of the connectome [48, 30] suggesting that the dense connectivity of cortical hubs, known as the “rich club”, gives rise to slow stable dynamics in core areas that include much of the neuronal circuitry pertaining to emotion and cognitive control. On the other hand, there are peripheral regions typically on the primary sensory cortex that create fast and unstable fluctuations [46]. These findings suggest that there is a correspondence of the slow time scales of the emotional highly connected nodes with the slow time scales of internal states (such as mood changes that bias learning and expectations according to the real life context [21], whereas peripheral regions’ fluctuations are related to the fast alterations of events in the external sensory apparatus. Potential structural changes located in these core hub-regions could destabilize the aforementioned slow dynamics and could contribute to the mood variations of BD patients [48]. In this work we have revisited a neural model of bipolar disorder based upon mutual inhibition of neuronal populations. This core hypothesis of the model, is supported by various examples in the neuropsychiatric literature of mutual inhibition of two neuronal networks. More specifically, two assumed mutually inhibiting neural circuits, that produce bistability, seem to be involved in the mechanism of the REM-nonREM transitions during sleep [42, 35, 29]. The dynamical consequences of mutual inhibition have also been studied in two-neuron models [69], while the effect of mutual inhibition supplemented with auto-inhibition has been investigated in a theoretical study of Reticular Thalamic oscillations [18]. Moreover, Ramirez-Mahaluf et al. [54, 53], after establishing the opposite functions and anticorrelation of dorsolateral prefrontal cortex (dlPFC) and ventral anterior cingulate cortex (vACC) during the performance of emotional and cognitive tasks, they propose a computational model that explains Major Depressive Disorder (MDD) symptoms and treatments. This model is based on the hypothesis of mutual disynaptic inhibition between the vACC and the dlPFC networks.

In summary, we were able to show that the complex oscillations obtained and reported by Goldbeter [28, 27] can be understood as MMOs due to a folded node, which we could provide analytical and numerical evidence of, once we found the appropriate timescale separation in the initial model. Furthermore, we explored the bifurcation structure of the model with respect to the parameter *K*_*f*1_ and with respect to both *K*_*f*1_ and *V*_*M*_. The former bifurcation analysis led to the identification of 3 distinct regimes of the system, namely that of depression, bipolar disorder and mania. The transitions between those regimes are characterised by the existence of MMOs that in parameter space are created by different bifurcations in each case. The latter bifurcation analysis on the other hand, led to the identification of the range of values of the two parameters for which there is an oscillatory regime (corresponding to bipolar disorder) and for which there is the possibility of the two distinct steady states (depression and mania) that were achievable for low and high values of *K*_*f*1_, respectively. Moreover, we show that the addition of small noise (of order *ε*) to the two fast variables of the system does not affect the MMO structure and we propose a minimal model consisting of two fast variables and one slow variable with a noise term that also produces MMOs which could correspond to the induction of the mixed bipolar states by random life events. Finally, we found numerical evidence of deterministic chaos produced by this model. For a specific value of the parameter *K*_*f*1_, possibly through a period-doubling cascade, we find by direct simulation a trajectory which seemingly follows a chaotic attractor. The broader significance of our work is that we introduce to the neuropsychiatric community some relatively new mathematical tools which enable the in-depth analysis of systems evolving on multiple timescales. This type of analysis and control could be applied to future more biologically plausible models of bipolar disorder. In particular, one could also derive information about the values of specific parameters from the regime that the system is in, through the bifurcation diagrams. More generally, multiple-timescale dynamical modeling, which is customary in a number of biological topics (such as neuronal and cardiac cells’ electrical activity, cell cycles, hormonal release, to name a few), should become a part of the toolbox of computational psychiatry [2, 59] and we provide a case study based on BD to demonstrate its efficiency.

Finally, among the questions that remain open are those regarding the biological meaning of the parameters of the system. In this study we focused on parameters *K*_*f*1_ and *V*_*M*_. As explained in Sections 3.1 and 3.2, these parameters directly govern the rate at which the variables of the model increase in time. Parameters *V*_*D*_ and *V*_*M*_ dictate the maximum rate of increase of the propensity to depression and mania respectively, which are correlated with the activation of the two opposing neuronal populations *D* and *M*, while *K*_*f*1_, and symmetrically *K*_*f*2_, measure the levels of *M* or *D* achieving 50% of the maximum rates of production of the intermediate factors *F*_*M*_ or *F*_*D*_, respectively. These parameters are likely affected by molecules used to treat bipolar disorders, such as antidepressants. The model parameters encompass a variety of processes controlling the electrophysiological activity of the neural network involved. These processes may range from the control of ionic conductances and synaptic function to the rates of synthesis, transport or degradation of hormones and neurotransmitters, as well as the affinity of the receptors that respond to these signals. At the molecular level, a more thorough physiological characterization of the model parameters awaits the identification of the molecular and cellular mechanisms involved in the origin of bipolar disorders.

The interest of the modeling approach is to pinpoint the type of plausible mechanisms that are capable of giving rise to the different modes of dynamic behavior associated with this neuropsychiatric disorder. The analysis that we present provides a detailed mathematical description of the transitions between these various modes of dynamic behavior, which may hopefully contribute to a better understanding of bipolar disorder and its control. Aspects of this topic that could be explored further are: 1) using a generalized methodology for dissecting systems with more than 2 fast and 2 slow variables and improving our numerical results, 2) studying how manipulating the ratio of the eigenvalues of the Jacobian matrix of the DRS could affect the number of smaller oscillations that we observe and the delay in the transition between the two different regimes, depression and mania. In a later stage of research, this could lead to the identification of a specific biological substrate that generates the observed oscillations as well as a better stratification of patients with respect to their individualized set of parameters. In any case, there is potential for further research as this approach for dissecting the dynamics of complex systems that exhibit this kind of periodic behavior seems very promising and calls for applications in diverse fields that range from bursting neuron models to mood oscillations in psychiatric patients.

## Acknowledgments

This work came into fruition during Efstathios Pavlidis’ internship in MathNeuro team, Inria, for the M.Sc. program: “Modeling for Neuronal and Cognitive Systems” (Mod4NeuCog) offered by the NeuroMod Institute and the Université Côte d’Azur (France0.

## A Appendix multiple-timescale analysis of the model

### A.1 Fast dynamics and fast subsystem

The full system of ODEs, written explicitly as a slow-fast system in fast-time formulation (timescale ratio parameter *ε* multiplying the right hand side of the slow variables):

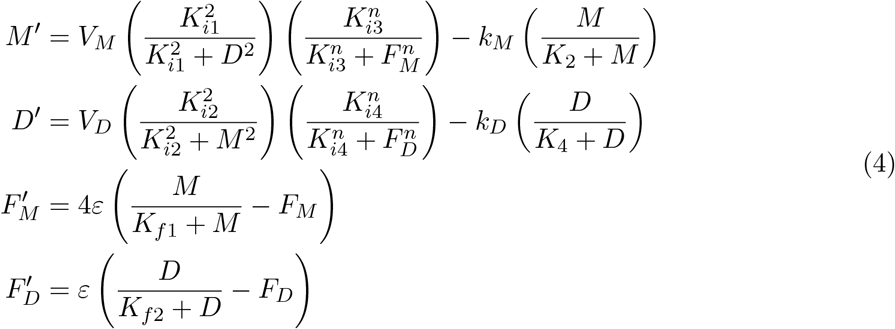

Taking the limit *ε* = 0 of the fast-time system yields the so-called *fast subsystem*, in which *F*_*M*_ and *F*_*D*_ have their dynamics frozen and are considered as parameters. The fast subsystem, that provides an approximation of the fast dynamics of the original system is:

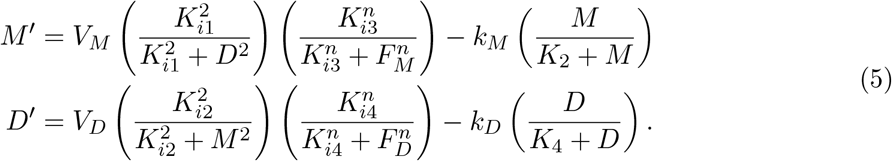

In order to find all the equilibria of the fast subsystem, we solve the equations D’=0, M’=0. The corresponding object is called the critical manifold of the system. It is a 2D surface (equilibria of fast subsystem and phase space of the slow subsystem) that we aim to plot in the 3D phase space. By performing algebraic manipulations to the previous equations, if we take *n* = 1 as in the papers by Goldbeter, we can solve for *F*_*M*_ :

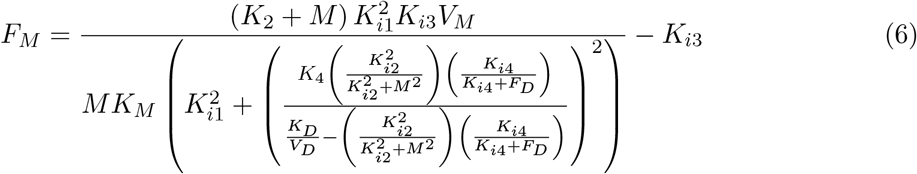

Plotting this formula for *F*_*M*_ in 3D using MATLAB yields the surface in figures 3 and 4

### A.2 Slow dynamics, Slow sub-system and Desingularised Reduced System (DRS)

We are interested in finding the limit of the slow segments of the MMO solution, or in other words what are the dynamics on the critical manifold, as the timescale ratio parameter *ε* tends to 0. The introduction of the so-called slow time *τ* = *tε* allows to put the system in its slow-time parametrization.

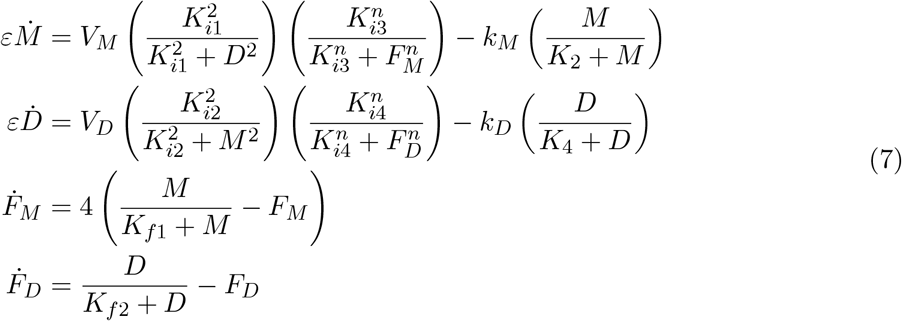

By taking the limit *ε* = 0 in the slow-time parametrization of the system, we obtain the *slow subsystem* that provides an approximation of the slow dynamics of the original system. The slow subsystem or reduced system (RS) is:

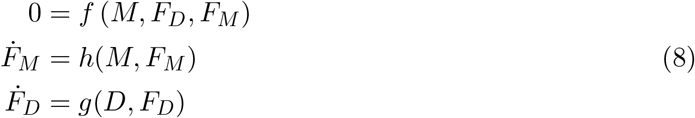

Note that the slow subsystem is a differential-algebraic system, that is, a system of two differential equations (for the original slow variables *F*_*M*_ and *F*_*D*_) constrained by an algebraic equation, which effectively corresponds to the critical manifold. Hence the critical manifold plays a key role in both subsystems: it is the set of equilibria of the fast subsystem, and it is the phase space of the slow subsystem. Since the algebraic constraint must be true for all time t, we can differentiate it with respect to t and we obtain:

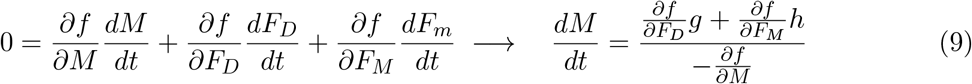

Therefore when 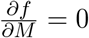 the RS is not defined. It is worth mentioning that the condition 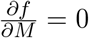 corresponds to the fold set of the critical manifold (S-shaped surface), locally formed by two curves (only one curve with two branches that meet at a cusp point) and it can be computed by continuing the fold bifurcation points of the fast subsystem in both *F*_*M*_ and *F*_*D*_. This is the reason to introduce an auxiliary system called the Desingularized Reduced System (DRS) obtained by rescaling time in the RS by a factor “−*df/dM* “. If we apply the convention that the flow of the RS and of the DRS should have the same direction on the attracting sheet of the critical manifold, which corresponds to the submanifold {*df/dM <* 0} and opposite direction on the repelling sheet of S, which corresponds to the submanifold {*df/dM >* 0}, then the only possible way to obtain the DRS from the RS is to introduce an auxiliary time *s* = −*tdf/dM* and we obtain the following Desingularized Reduced System (DRS):

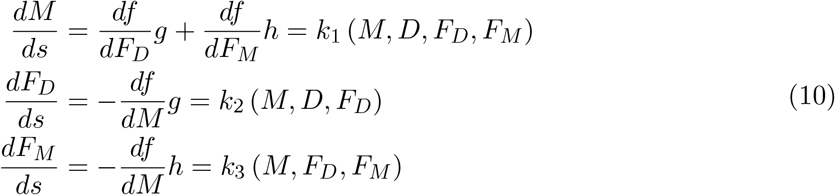

With the process of desingularization essentially, we project the points of the critical manifold (a surface) onto a plane, which removes the singularity along the fold set, and we also change the orientation of the flow along the repelling sheet. Note that the RS is a dynamical system constrained to evolve on a surface (the critical manifold), therefore it can be described with only two equations. Since we have an explicit formula for *F*_*M*_ that depends only on M and *F*_*D*_, we can eliminate the equation for *dFm/ds*. The DRS becomes:

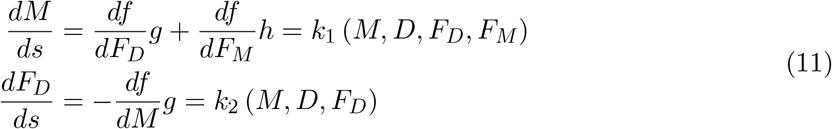

For the DRS to have an equilibrium two conditions must be satisfied: 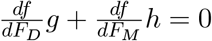 and 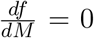 or *g* = 0. The equilibria of the DRS satisfying 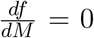 are precisely the ones that have been created by the process of desingularization allowing to pass from the RS to the DRS. Therefore, they are of interest to us since we want to understand the dynamics along the fold set of the critical manifold, where the RS is not defined. Such equilibria of the DRS are called folded equilibria for the RS. The DRS, in other words, offers a way to understand the slow flow up to the fold curve via the existence of folded equilibria, which are true equilibria of the DRS located on the fold curve. The fact that the right-hand side of the M-equation of the RS has a denominator going to 0 on the fold curve of the critical manifold makes the RS a priori undefined along this curve. However, when the term 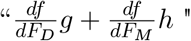 also has a zero of the same order as 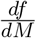, then 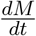 is well defined and so is the RS. Therefore, even though the RS is undefined at most points on the fold curve, there are specific points where it is defined, and these are folded singularities. The conditions for a folded singularity **p** are the following:

1. *F* (**p**) = 0 (critical manifold)
2. 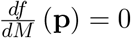 (fold curve)
3. 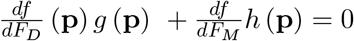

It is worth noting that folded singularities are not equilibria of the RS but important points that allow a dynamical passage from the attracting onto the repelling sheet of the critical manifold. This corresponds to a canard type dynamic in the singular limit *ε* = 0, and therefore the corresponding solutions of the RS are called singular canards [17]. To find the type of the folded singularity of the DRS we can compute the eigenvalues of its Jacobian matrix **A**:

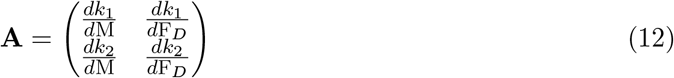

In order to find the eigenvalues for the fold point **p**, we must solve det(**A** − *λ***I**) = 0, with **A** evaluated at the point **p**. If *s*_1_, *s*_2_ are the eigenvalues of the Jacobian Matrix A evaluated at the folded singularity **p**, as an equilibrium of the DRS, then **p** is a folded node if *s*_1_ *<* 0, *s*_2_ *<* 0 and *s*_1_, *s*_2_ ∈ ℝ.

### A.3 Robustness of MMOs to noise

We add low-amplitude noise terms, of the order of *ε*, to the two fast variables:

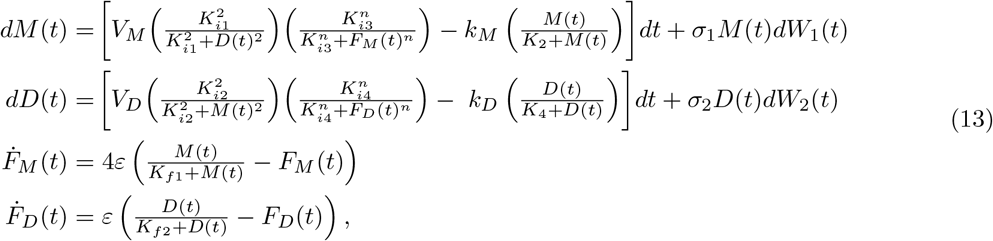

where *W*_*i*_(*t*) are independent standard Wiener processes (also independent of the initial conditions when the latter are stochastic), and where the structure of the drift terms (i.e. the terms in “*dt*”) and of the diffusion terms (i.e. the term in “*dW*_*i*_(*t*)”) ensures that *M* (*t*) and *D*(*t*) remain positive almost surely for all *t* ≥ 0: indeed, for in *M* (*t*) = 0 (resp. *D*(*t*) = 0) the noise term is null and the drift term (i.e. the term in “*dt*”) is positive in the corresponding equation.

### A.4 Noise induced MMOs

Removing the variable *F*_*D*_ and adding a noise term in the other slow variable *F*_*M*_ :

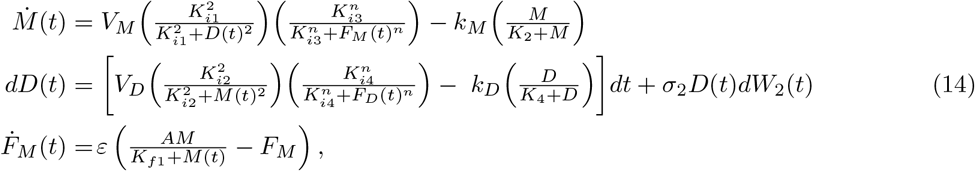

where a standard Wiener process with a diffusion term *σ*_2_ *D*(*t*) is added to the dynamics of *D*, as before, when *D*(*t*) = 0, the diffusion term is null and the drift term is nonnegative so that *D*(*t*) stays nonnegative for all *t* ≥ 0, almost surely. The thought process behind the construction of the 3D stochastic model is the following: the 2D system with only of *M* and *D* variables, with *F*_*M*_ and *F*_*D*_ being parameters, as described by A. Goldbeter [27, 28] exhibits bistability between equilibria (has an S-shaped curve of equilibria) which will allow for relaxation oscillations when slow variable(s) are added (e.g. *F*_*M*_). When slow dynamics is added for *F*_*M*_, while *F*_*D*_ is still a parameter, the resulting 3D system displays bistability between large-amplitude relaxation cycles and equilibria, when *F*_*D*_ is statically varied. The system has 2 Hopf bifurcations which are both subcritical and the initially unstable branches of limit cycles become stable through saddle-node bifurcations of cycles. This system has two zones of bistability (in between the Hopf points and the saddle-node of cycles point) between equilibria and limit cycles. Then, as the noise term *σ*_2_*D*(*t*)*dW*_2_(*t*) is introduced to the system, its effect is essentially to make the system “jitter” in the zone of bistability. Because the branch of cycles without noise grows sharply (canards), this is the reason why the noisy system exhibits noise-induced MMOs; see e.g. [60] for an example of analysis of such noisy complex oscillations.

## Footnotes

1 https://senselab.med.yale.edu/ModelDB/

